# The Atomic-Level Physiochemical Determinants of T Cell Receptor Dissociation Kinetics

**DOI:** 10.1101/2021.10.25.465739

**Authors:** Zachary A. Rollins, Jun Huang, Ilias Tagkopoulos, Roland Faller, Steven C. George

**Affiliations:** Department of Biomedical Engineering, University of California, Davis, Davis, California; Department of Chemical Engineering, University of California, Davis, Davis, California; Department of Computer Science, University of California, Davis, Davis, California; Pritzker School of Molecular Engineering, University of Chicago, Chicago, IL

**Keywords:** T cell receptor, peptide major histocompatibility complex, immunogenicity, steered molecular dynamics, machine learning

## Abstract

The rational design of T Cell Receptors (TCRs) for immunotherapy has stagnated due to a limited understanding of the dynamic physiochemical features of the TCR that elicit an immunogenic response. The physiochemical features of the TCR-peptide major histocompatibility complex (pMHC) bond dictate bond lifetime which, in turn, correlates with immunogenicity. Here, we: i) characterize the force-dependent dissociation kinetics of the bond between a TCR and a set of pMHC ligands using Steered Molecular Dynamics (SMD); and ii) implement a machine learning algorithm to identify which physiochemical features of the TCR govern dissociation kinetics. Our results demonstrate that the total number of hydrogen bonds between the CDR2β-MHCα(β), CDR1α-Peptide, and CDR3β-Peptide are critical features that determine bond lifetime. We propose that amino acid substitutions to these hypervariable regions of the TCR can efficiently manipulate immunogenicity and thus be used in the rational design of TCRs for immunotherapy.

## INTRODUCTION

T cell-based immunotherapies (e.g., chimeric antigen receptor-T, or CAR-T; and TCR-engineered-T, or TCR-T) have provided transformative therapeutic responses in a small subset of cancers and patients^1-5^; however, progress in solid tumors has been agonizingly slow. For example, CAR-T cells require an antigen on the tumor cell surface, but the majority (∼85%) of identified neoantigens are intracellular^6^ and thus are immunogenic only when a representative fragment is presented on the cell surface in a peptide-major histocompatibility complex (i.e., pMHC). Although TCR-T therapy is MHC-restricted, this approach can target intracellular antigens, and the remarkable sensitivity of a TCR to recognize a single pMHC molecule^7^ provides an additional strategic advantage. Nonetheless, identifying neoepitopes, matching these with immunogenic TCRs, and minimizing off-target effects remain significant challenges to implementation of these therapies^8^.

Recent reports demonstrate that single-cell sequencing and machine learning technologies can identify patient- and tumor-specific neoepitopes^9, 10^. However, identification of partner TCRs remains challenging, despite the fact that tumor-specific T cells can be found in the peripheral blood^11, 12^. The human immune system generates tumor-specific T cells in a process that begins with random V(D)J recombination to create the hypervariable regions of the TCRα and β chains. While this process generates a stunningly large number of *possible* TCRs (>10^20^-10^61^)^13, 14^, including 10^6^-10^8^ in the peripheral blood, it is inherently inefficient and does not necessarily produce a TCR with appropriate immunogenicity for a given tumor^15^. Alternate strategies of TCR identification have also fallen short; for example, TCR affinity enhancement can lead to a loss of TCR specificity^16, 17^ and does not always determine immunogenicity^18^.

Computational techniques such as steered molecular dynamics (SMD) and machine learning may enable the creation of highly immunogenic, tumor-specific TCRs through rapid and efficient screening of the vast number of possible TCRs. The success of these techniques depends on accurate *in vitro* predictions of T cell immunogenicity, a goal that remains elusive. Quantitative descriptors of the TCR-pMHC bond identified in previous studies do not consistently correlate with immunogenicity^18-21^. The majority of these studies measured equilibrium parameters of the TCR-pMHC bond, which do not account for the mechanical forces on the TCR-pMHC bond present *in vivo*. Recent studies using DNA-based tension probes have estimated this force at ∼10-20 pN^22, 23^, and subsequent studies demonstrate that dissociation kinetics (i.e., bond lifetime) of the TCR-pMHC bond at this physiologic force can predict immunogenicity^24-31^. These correlations are consistent across species, TCR-pMHC pairs, and experimental systems^24-31^.

Here, we seek to discern the atomic-level physiochemical features that determine the TCR-pMHC bond lifetime under force (i.e., characterize the TCR-pMHC’s force-dependent dissociation kinetics). As a first attempt to manipulate the bond lifetime of the TCR-pMHC over a wide range, we characterized the force-dependent dissociation kinetics of a single TCR (with a known crystal structure) to 17 possible pMHCs using steered molecular dynamics (SMD). Then, we used machine learning to identify the physiochemical features and the specific regions of the TCR regulating bond lifetime. Our results demonstrate that the total number of hydrogen bonds (H-bonds) between the CDR2β-MHCα(β), CDR1α-Peptide, and CDR3β-Peptide are critical features that determine bond lifetime. This finding may inform the rational design of TCRs for TCR-T cell therapy, and provide a path forward to create more advanced and predictive maching learning algorithms.

## METHODS

### Molecular Dynamics Setup

The crystal structure of the human DMF5 TCR complexed with agonist pMHC MART1-HLA-A2 (PDB code: 3QDJ)^32^ was the initial structure for all simulations (**Figure 1A**). To generate the 17 TCR-pMHC pairs, amino acid substitutions were made to the MART1 peptide (AAGIGILTV) using the Mutagenesis plugin on Pymol Molecular Graphics System (Schrödinger, New York, New York). Interfacial substructures (**Figure 1B**) were defined by sequential residues from the corresponding chains: TCRα (CDR1α: 24-32, CDR2α: 50-55, CDR3α: 89-99), TCRβ (CDR1β: 25-31, CDR2β: 51-58, CDR3β: 92-103), MHCα (MHCα(β): 50-85, MHCα(α): 138-179), and peptide (1-9). To determine protonation states, pKa values were calculated using propka3.1^33, 34^ and residues were considered deprotonated in Gromacs^35^ if pKa values were below the physiological pH 7.4. The resulting systems were solvated in rectangular water boxes using the TIP3P water model^36^ large enough to satisfy the minimum image convention. Na^+^ and Cl^-^ ions were added to neutralize protein charge and reach physiologic salt concentration of ∼150 mM. All simulations were performed with Gromacs 2019.1^35^ using the CHARM 22 plus CMAP force field for proteins^37^ and orthorhombic periodic boundary conditions. All simulations were in full atomistic detail.

**Figure 1:**
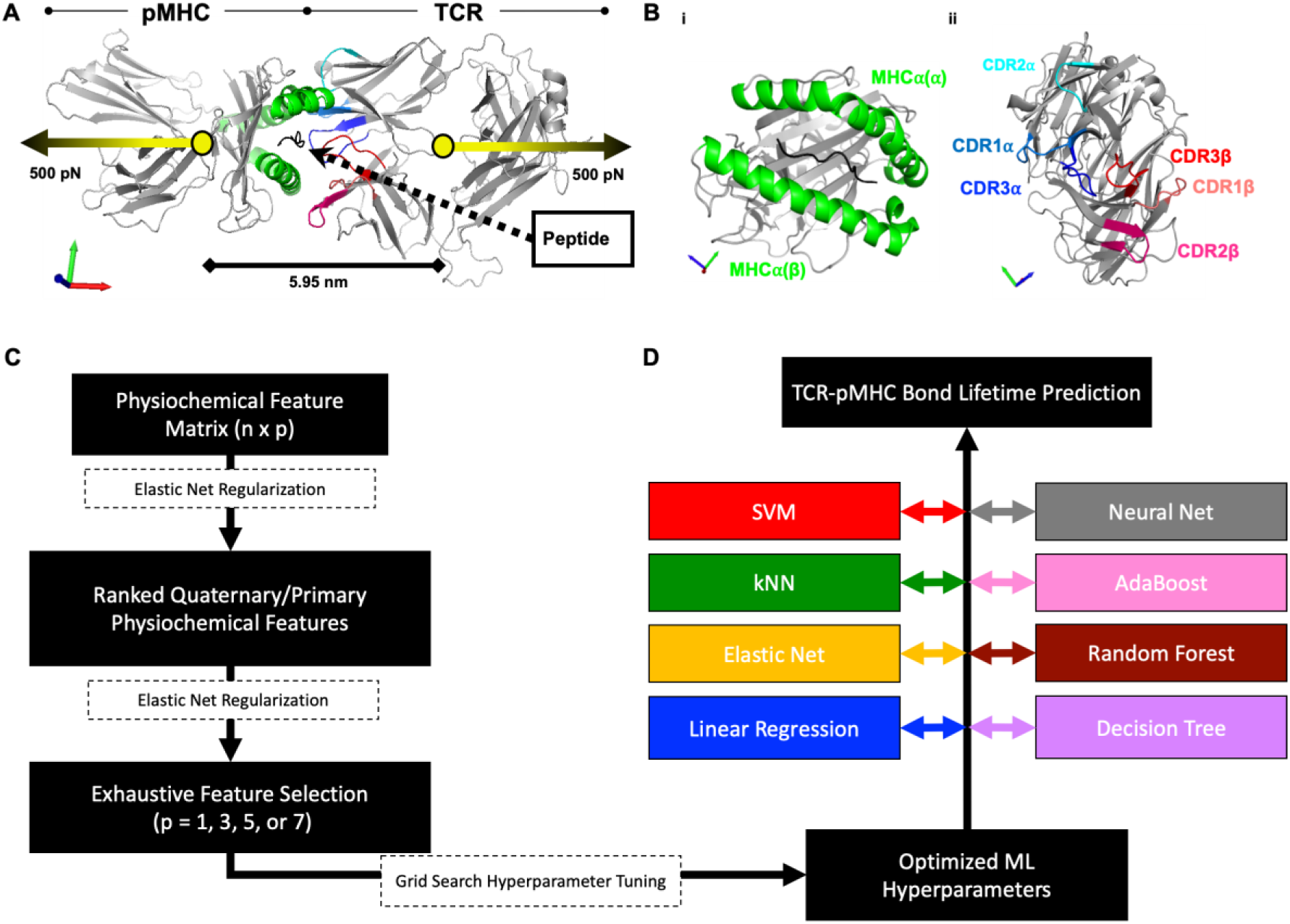
Steered Molecular Dynamics (SMD) simulations and machine learning algorithms were used to identify the physiochemical features that predict TCR-pMHC bond lifetime. **(A)** Starting structure for SMD of TCR and pMHC (shown at the top with black lines and circle arrowheads). The location/direction of pulling are depicted with yellow circles/arrows, respectively; the black scale bar with diamond arrowheads denotes the locality of distance between center of masses. The non-interacting bodies of the TCR and pMHC are colored in gray. Axis directions are indicated in left corner (red: +x-direction, blue: +y-direction, and green: +z-direction). **(B)** The primary interfacial substructures: **(i)** MHCα(α) & MHCα(β) = green, Epitope=black; and **(ii)** TCR CDR1α = light blue, TCR CDR2α = cyan, TCR CDR2α = dark blue, TCR CDR1β = salmon, TCR CDR2β = light red, and TCR CDR3β = red. **(C)** A two-layer Elastic Net-Exhaustive Feature Selection algorithm (dashed boxes) was used to obtain ranked and reduced feature sets. **(D)** Selected features were used to tune hyperparameters (dashed box) for each machine learning model (Linear Regression = blue, Elastic Net = orange, k-Nearest Neighbors = green, Support Vector Machines = red, Decision Tree = purple, Random Forest = brown, AdaBoost = pink, Neural Net = gray).

### Energy Minimization and Equilibration

Generating equilibrated starting structures for the Steered Molecular Dynamics simulations required four steps: (1) Steepest descent energy minimization to ensure correct geometry and the absence of steric clashes; (2) 100 ps simulation in the constant volume (NVT) ensemble to bring atoms to correct kinetic energies, while maintaining temperature at 310 K by coupling all protein and non-protein atoms to separate baths using the velocity rescale thermostat with a 0.1 ps time constant^38^; (3) 100 ps simulation in the constant pressure (NPT) ensemble using Berendsen pressure coupling^38^ and a 2.0 ps time constant to maintain isotropic pressure at 1.0 bar; and (4) Production MD simulations conducted for 50-150 ns with no restraints. The protein structures were evaluated every 50 ns to determine if all protein chains were equilibrated by root mean square deviation. To ensure true NPT ensemble sampling during 100 ns production runs, the Nose-Hoover thermostat^39^ and Parrinello-Rahman barostat^40^ were used to maintain temperature and pressure, respectively. Time constants were 2.0 and 1.0 ps for pressure and temperature coupling, respectively, utilizing the isothermal compressibility of water, 4.5^-5^ bar^-1^. Box size for equilibration was 10.627 × 7.973 x.10.685 nm^3^ with ∼ 48,000 water molecules, ∼300 ions, and ∼157,000 total atoms. All simulation steps used the Particle Ewald Mesh algorithm^41, 42^ for long-range electrostatic calculations with cubic interpolation and 0.12 nm maximum grid spacing. Short-range non-bonded interactions were cut off at 1.2 nm using the Verlet cutoff-scheme and all bond lengths were constrained using LINCS algorithm^43^. The leap-frog algorithm was used for integrating equations of motion with 2 fs time steps. After the preparation runs, three independent MD configurations for each peptide mutant were extracted and used as the three starting points for steered molecular dynamics simulations.

### Steered Molecular Dynamics (SMD)

The full TCR-pMHC complex structure was extracted from the preparation run for each peptide mutant to generate three SMD starting configurations. The x-axis of these protein complexes was aligned along the *x*-axis and solvated in rectangular water boxes with dimensions 30 × 9.972 × 12.685 nm^3^. Solvent was again represented by the TIP3P water model and Na ^+^ and Cl^-^ ions were added to neutralize protein charge and reach physiologic salt concentration of ∼150 mM. This resulted in ∼120,000 water molecules, ∼700 ions, and ∼370,000 total atoms. All Gromacs structure files are uploaded to the Dryad repository for the exact atomic specifications. Before pulling, all systems underwent (1) energy minimization; (2) 100 ps NVT; and (3) and 100 ps NPT to remove high energy contacts without disturbing the configurations. During pull, the Nose-Hoover thermostat and Parrinello-Rahman barostat were used to maintain temperature and pressure. 500 pN linear potential was applied to the center of mass (COM) of the TCR and pMHC in the *x*-direction and simulations continued until distance between COMs reached 0.49 times the box size in *x*-direction (**Figure 1A**). The COM was chosen as the site of applied force because pulling from the TCR and MHC termini resulted in artificial unfolding (not shown). All simulation trajectories and selected frames were visualized using the Pymol Molecular Graphics System (Schrödinger, New York, New York).

### Physiochemical Descriptors and Data Analysis

Physiochemical descriptors were evaluated by defining Gromacs index groups (gmx make_ndx) and using Gromacs-suite analysis tools (i.e., gmx hbond, gmx rms, gmx rmsf, gmx sasa, gmx gyrate, gmx distance). Data analyses were performed by standard python packages for data handling and visualization (i.e., numpy^44^, pandas^45^, seaborn^46^, matplotlib^47^, statistics^48^, and GromacsWrapper^49^), and custom python scripts. Random mutants were generated with a custom python script compatible with Pymol using the random python package and selecting a random location and amino acid to mutate the peptide. The machine learning algorithms were developed using the sklearn package^50, 51^ and exhaustive feature selection was performed using mlxtend package^52^. The geometry of a Lennard-Jones contact (LJ-contact) is defined as a distance less than 0.35 nm between atoms. The L1 peptide bond lifetime was an outlier (z-score = 3.65 > 3). To reduce the effects of the outlier on the dataset, median absolute error was selected as the scoring criterion and L1 was excluded from correlation coefficient calculations. The mean absolute error represents the arithmetic average of median absolute error from repeated three-fold cross validation. The Pearson correlation coefficient (r_p_) and Spearman rank correlation coefficients (r_s_) were calculated using the correlation method in the pandas python package. Akaike and Bayesian Information Criterion (AIC and BIC) were calculated from the standard deviation of repeated three-fold cross validation of the best machine learning algorithm selected from the hyperparameter grid search. Statistical significance was determined by performing a one-tailed student’s t-test (p<0.05) for each machine learning algorithm across feature sets. Custom scripts relevant to mutant generation, feature selection, machine learning, and the production of figures have been made available on a GitHub repository: https://github.com/zrollins/TCR.ai.git.

### Feature Selection and Machine Learning Algorithms

Features were ranked and reduced utilizing a two-layer Elastic Net – Exhaustive Search algorithm (**Figure 1C**). First, Elastic Net Regularization^53^ was used with all physiochemical features and a grid search was performed to optimize hyperparameters. The optimized hyperparameters were implemented into the Exhaustive Feature Selector^52^ and the best individual features were ranked by repeated (n_repeats=3) threefold cross-validation. The top ten features were ranked by mean absolute error and feature combinations were exhaustively searched, utilizing Elastic Net Regularization, to determine the best combinations of 3, 5, and 7 features (**Figure 1C**). The best feature combinations were selected by mean absolute error arithmetically averaged over the cross-validation. These feature combinations were then implemented into several machine learning algorithms to determine the most predictive model of bond lifetime (**Figure 1D**)^50, 51^. The machine learning algorithm hyperparameter optimization was performed on a high performance compute cluster at the University of California, Davis College of Engineering and the best model for each feature set was scored on absolute error and ranked by the arithmetic average of repeated threefold cross-validation (i.e., n_splits=3, n_repeats=3, random_state=1). Detailed documentation regarding the cross validation and hyperparameter optimization of two-layer Elastic Net – Exhaustive Search feature selection and machine learning predictions are provided in the supporting information. In addition, this dataset has been made freely available on the GitHub repository.

## RESULTS

### Bond lifetime

As the starting point to simulate the force-dependent dissociation kinetics of 17 TCR-pMHC pairs using SMD, we used the previously reported crystal structure (PDB ID: 3QDJ)^32^ of the DMF5 TCR (from a melanoma patient) bound to the MART1 peptide (AAGIGILTV)-MHC complex (**Figure 1A**). We then replaced the MART1 peptide with 16 different peptides (**Figure 1—supplement 1**) for a total of 17 TCR-pMHC pairs. Ten peptides were chosen from a set of known pMHCs^54, 55^ and 7 were generated through random point mutation of the MART1 peptide. For these 17 TCR-pMHC pairs, the mean bond lifetime in the SMD simulations was 5400 ± 1700 picoseconds (**Figure 2**).

**Figure 2.**
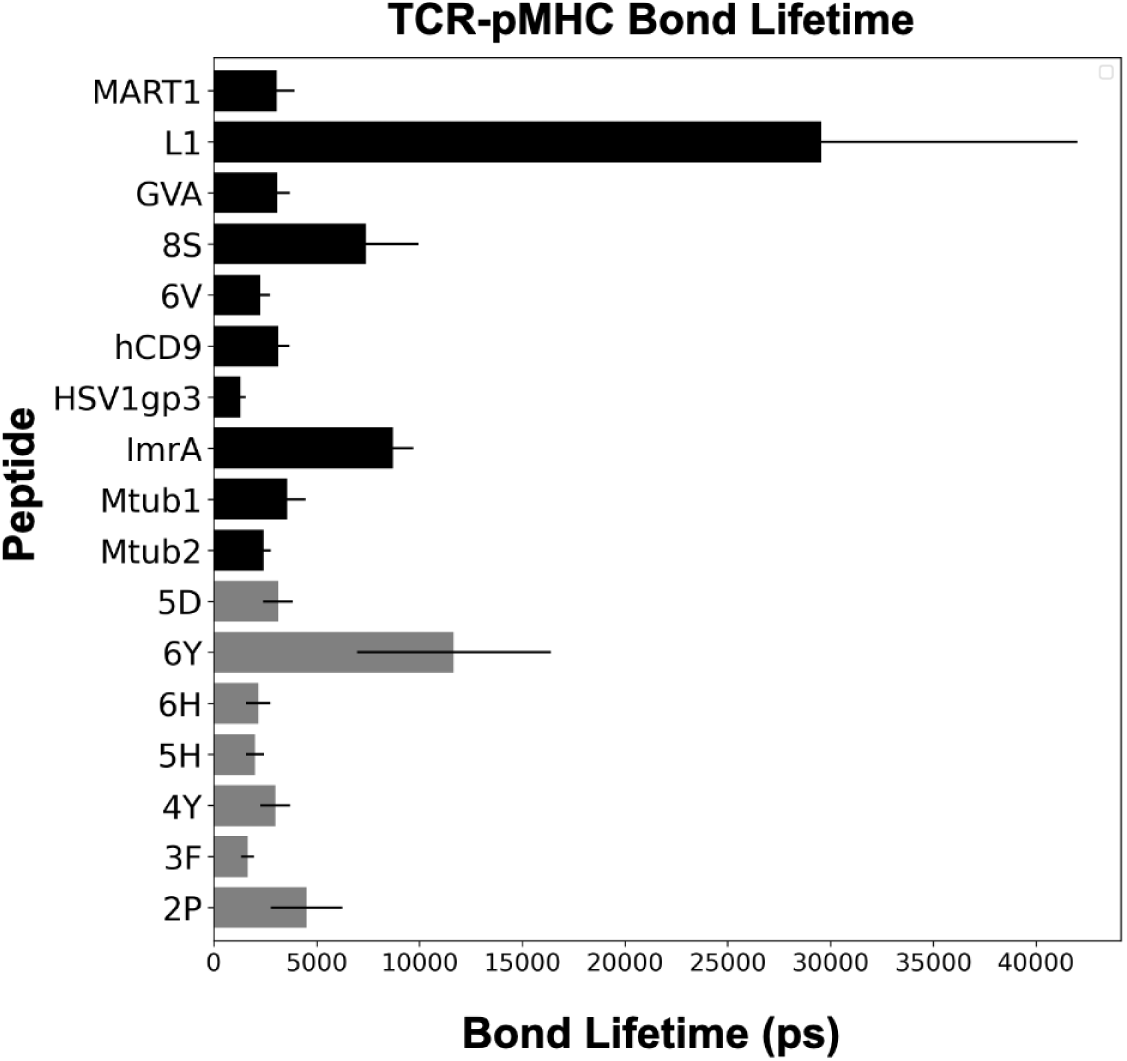
Mean TCR-pMHC bond lifetime for 17 different peptides. Using Steered Molecular Dynamics (SMD), we applied a constant force of 500 pN at the center of mass for the TCR and pMHC and estimated the mean bond lifetime for 17 different peptides. Known peptides and those with random point mutations are denoted with black and gray bars, respectively. Each TCR-pMHC was pulled apart 3 times using different equilibrated structures.

### Physiochemical features of the TCR-pMHC

Next, we identified two sets of physiochemical features which, at distinct resolution levels, describe the TCR-pMHC bond during the SMD simulation. The first set characterizes physiochemical features of the entire TCR-pMHC interaction (e.g., total H-bonds between the TCR and pMHC). This characterization provides an overall assessment of the physiochemical features that might impact bond lifetime and is consistent with the quaternary structure of globular proteins. We considered features likely to impact dissociation kinetics and thus included H-bonds^56^, LJ-contacts^57^, distance between the TCR and pMHC^58, 59^, solvent accessible surface area (SASA)^60^, root mean square fluctuations (RMSF)^61^, and the gyration tensor of the TCR and pMHC. This approach resulted in 18 features for the first set, and we dubbed these quaternary features (**Figure 1—supplement 2**).

An understanding of the physiochemical features that regulate dissociation kinetics of the global TCR-pMHC bond provides an overall assessment of which physiochemical features regulate bond lifetime. However, this approach does not identify the sub-regions of the TCR-pMHC bond that regulate bond lifetime and thus are suitable targets for rational design of TCRs. The hypervariable regions of the TCR can be divided into 3 complementarity determining regions (CDRs) on the α and β chain, respectively. Within the MHC, the peptide is surrounded by α-helices which also interact with the nearby chains of the TCR (**Figure 1B**). These MHCα-helices are located on the MHCα chain and these substructures are defined by their interaction with the TCRα and β chain, respectively (i.e., MHCα(α) and MHCα(β)). These TCR CDRs and MHCα-helices form an interface with the peptide antigen – the variable in this study – and based on their physical location are likely to influence TCR-pMHC bond lifetime. Hence, we also identified a second set of features focused on the interface between the TCR and the pMHC (e.g., CDR3α loop of the TCR and the MHCα(β) chain, **Figure 1B**). This higher level of resolution is consistent with the secondary structures (e.g., α-helices) of a protein. Again, we considered features that are likely to affect dissociation kinetics and thus included H-bonds, LJ-contacts, distance between the sub-regions, SASA, RMSF, and the gyration tensor of the sub-regions. From these considerations, we identified 79 secondary features (**Figure 1—supplement 3**) that could potentially impact dissociation kinetics. The quaternary and secondary features were further categorized into chemical – such as H-bonds and LJ-Contacts – and physical – including RMSF, SASA, and the gyration tensor – interaction parameters.

### TCR-pMHC Bond Lifetime Prediction using Quaternary Physiochemical Features

To examine how quaternary physiochemical features influence TCR-pMHC bond dissociation kinetics, we ranked the top ten quaternary features after an Elastic Net grid search for each individual feature (**Figure 3A**). The scoring criterion was mean absolute error of bond lifetime in picoseconds. After Elastic Net grid search, chemical interaction features, in particular Total LJ-contacts and Total H-bonds, were the most predictive (**Figure 3A**); in particular, the total number of unique LJ-Contacts between TCR and pMHC had the smallest mean absolute error. In addition, the total LJ-Contacts had the highest Pearson and Spearman correlation coefficients (**Figure 3—supplement 1, Figure 3—supplement 3**).

**Figure 3:**
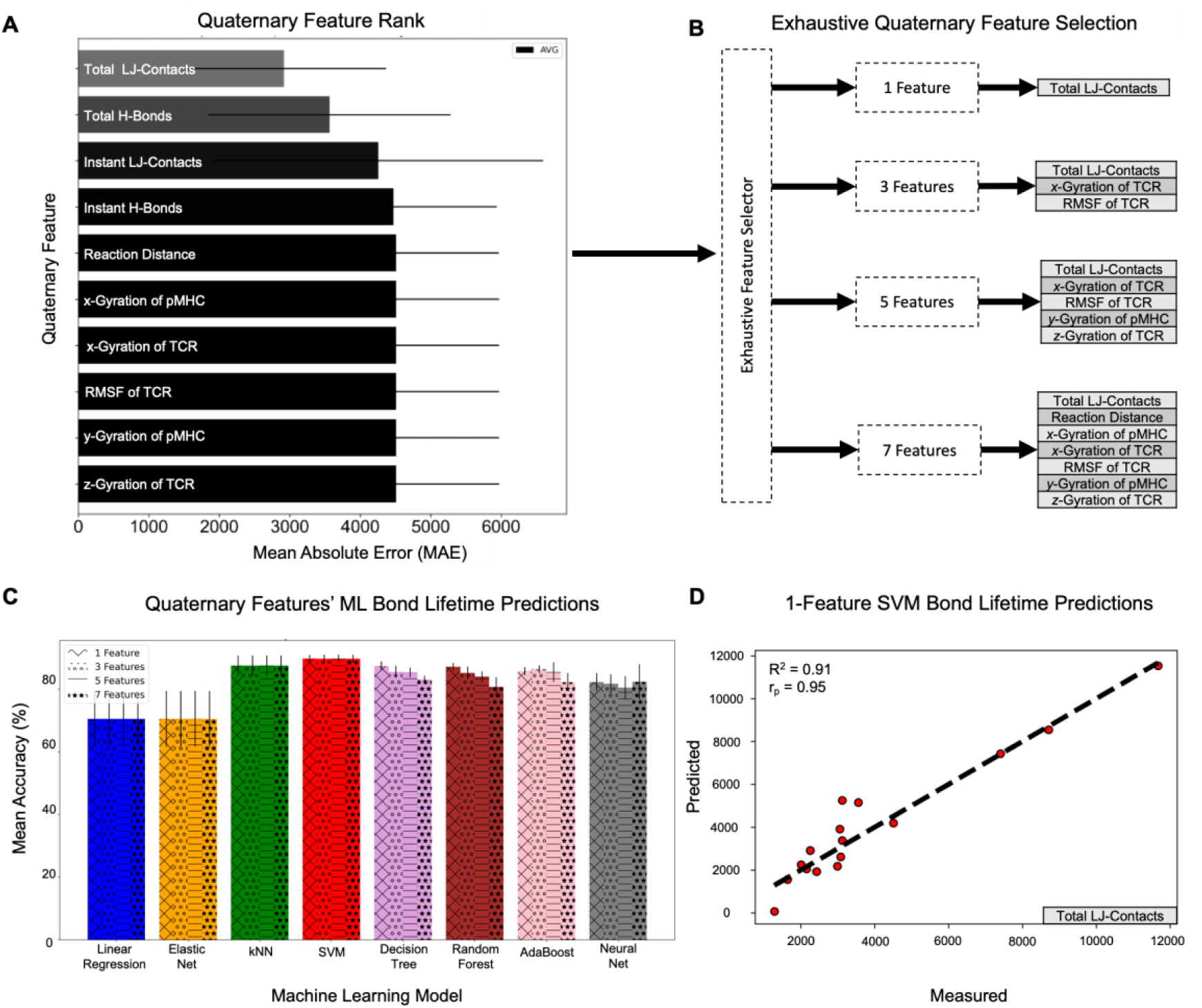
Quaternary Feature Selection and Bond Lifetime Predictions. **(A)** Mean absolute test error from elastic net regularization was used to select the top ten quaternary features. Errors represent the best test set standard deviation from repeated threefold cross-validation. **(B)** According to an exhaustive search, the best feature sets (i.e., p = 1, 3, 5, and 7) to predict bond lifetime. **(C)** The mean accuracies of bond lifetime prediction for all feature sets in (B) and machine learning models after hyperparameter tuning (Linear Regression = blue, Elastic Net = orange, k-Nearest Neighbors = green, Support Vector Machines = red, Decision Tree = purple, Random Forest = brown, AdaBoost = pink, Neural Net = gray). Errors represent the best test set standard error from repeated threefold cross-validation. The machine learning model standard error from cross-validation (n=9) was statistically compared for increasing feature sets by a one-tailed student’s t-test: #p<0.10, *p<0.05, **p<0.01. **(D)** The scatter plot of predicted and measured bond lifetimes from the selected one-feature Support Vector Machines algorithm with the coefficient of determination (top left), the Pearson correlation coefficient (top left), and the feature set (bottom right).

We next explored whether a combination of quaternary physiochemical features would improve predictions of bond lifetime. To accomplish this, we applied a regularized regression method (Elastic Net; see **Methods**) as a filter to identify predictive feature sets. To avoid overfitting^62-64^, feature sets were reduced utilizing an Elastic Net^53^ – Exhaustive Search^52^ algorithm (**Figure 1C**) to determine the best combinations of 3, 5, and 7 features. Using these combinations, we then trained and tested 8 different machine learning algorithms to estimate TCR-pMHC bond lifetime **(Figure 1D**)^50, 51^. Although physical quaternary features were selected in this exhaustive search (**Figure 3B**), these did not significantly improve the predictive power of the machine learning models (**Figure 3C**). This finding holds for all machine learning algorithms, as determined by the lack of statistically significant increase in mean accuracy or decrease in information criteria scores (Akaike and Bayesian Information Criteria) with increasing model complexity (**Figure 3—supplement 2, Figure 3—supplement 4**).

The best feature combination and machine learning model was chosen based on the lowest error and standard deviation from repeated three-fold cross-validation. Our results demonstrated that a feature set of only LJ-Contacts combined with a Support Vector Machines is best at predicting bond lifetime (**Figure 3D**). The mean absolute error using Support Vector Machines was 560 ± 200 picoseconds producing an accuracy of 90.0 ± 3.7% (i.e., 1-560/5400).

### TCR-pMHC Bond Lifetime Prediction Using Secondary Physiochemical Features

Analogous to our strategy to assess quaternary features of the TCR-pMHC, we examined secondary features. We ranked the top ten secondary features after an Elastic Net grid search for each individual feature (**Figure 4A**). The total number of unique H-bonds between CDR2β -MHCα(β) generated the smallest mean absolute error (**Figure 4A**). In addition, the top three features had the highest Pearson and Spearman correlation coefficients (**Figure 4—supplement 1, Figure 3—supplement 3**).

**Figure 4.**
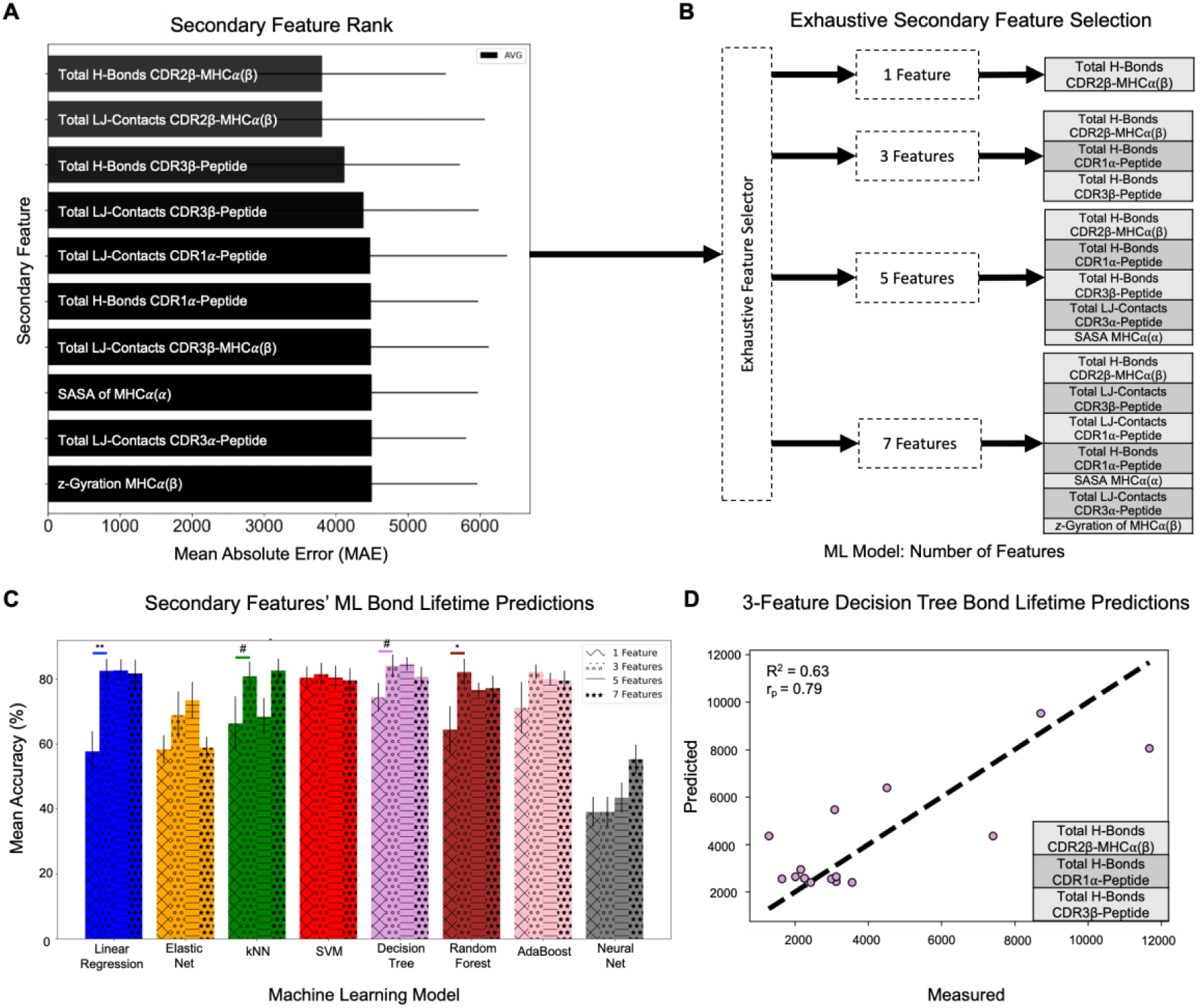
Secondary Feature Selection and Bond Lifetime Predictions. (**A**) Mean absolute test error from elastic net regularization was used to select the top ten secondary features. Errors represent the best test set standard deviation from repeated threefold cross-validation. (**B**) According to an exhaustive search, the best feature sets (i.e., p = 1, 3, 5, and 7) to predict bond lifetime. **(C)** The mean accuracies of bond lifetime prediction for all feature sets in (B) and machine learning models after hyperparameter tuning (Linear Regression = blue, Elastic Net = orange, k-Nearest Neighbors = green, Support Vector Machines = red, Decision Tree = purple, Random Forest = brown, AdaBoost = pink, Neural Net = gray). Errors represent the best test set standard error from repeated threefold cross-validation. The machine learning model standard error from cross-validation (n=9) was statistically compared for increasing feature sets by a one-tailed student’s t-test: #p<0.10, *p<0.05, **p<0.01. **(D)** The scatter plot of predicted and measured bond lifetimes from the selected 3-feature Decision Tree algorithm with the coefficient of determination (top left), the Pearson correlation coefficient (top left), and the feature set (bottom right).

We explored whether a combination of secondary physiochemical features would improve the prediction of bond lifetime. Following the same algorithm as for quaternary features, we applied an Elastic Net^53^ – Exhaustive Search^52^ algorithm (**Figure 1D**) to identify the best combinations of 3, 5, and 7 secondary features; cross-validated 8 machine learning models with these feature combinations; and selected the best feature combination and machine learning model based on error, standard deviation, and information criteria. Interestingly, the best 3 feature combination (CDR2β -MHCα(β), CDR1α-Peptide, and CDR3β-Peptide) selected by exhaustive search (**Figure 4B**) did not correspond to the top three individual features selected by Elastic Net rank (**Figure 4A)** or correlation coefficients (**Figure 3— supplement 3**). Compared to the single best feature, the best 3-feature combination statistically improved bond lifetime predictions for Linear Regression, k-Nearest Neighbors, Decision Tree, and Random Forest machine learning algorithms (**Figure 4C**). Increases in mean accuracy were not statistically significant beyond 3 features (**Figure 4C, Figure 4— supplement 3**). Moreover, these algorithms reduced information criteria scores (Akaike and Bayesian Information Criteria) when increasing from 1 to 3 features, whereas the Elastic Net, Support Vector Machines, and Neural Net algorithms increased both AIC & BIC (**Figure 4— supplement 2**). These results indicate that, among the secondary features and machine learning algorithms tested, a 3-feature combination utilizing a Decision Tree provides the most accurate prediction of bond lifetime (**Figure 4D**). The absolute error using the Decision Tree was 870 ± 570 picoseconds (**Figure 4—supplement 3**), or an accuracy of 84 ± 10%. In addition, this Decision Tree prediction by the best 3 feature combination exceeded the Pearson correlation coefficient of the individual features (**Figure 4D, Figure 4—supplement 2**).

## DISCUSSION

T cell-based immunotherapies, such as TCR-engineered-T cells, provide exciting potential to treat a wide range of cancers, including solid tumors. However, this potential has not been reached, due, in part, to the inability to rapidly and efficiently explore the vast TCR space to identify optimal tumor-specific TCRs. Experimental methods to design and test potential TCRs are expensive and slow, thus hindering throughput. In contrast, computational algorithms that utilize machine learning have enormous potential to rapidly interrogate the TCR space and identify a small number of candidates for more efficient experimental testing. We tested this premise using SMD to create a small database of TCR-pMHC bond lifetimes, then created machine learning algorithms to predict bond lifetime based on quaternary and secondary features of the TCR-pMHC bond. Using the quaternary features, we found that total LJ-contacts could predict bond lifetime with 90% accuracy. More importantly, we also found that we could predict bond lifetime with an accuracy of 84% using only the total H-bonds between three subregions of the TCR-pMHC: CDR2β-MHCα(β), CDR1α-Peptide, and CDR3β-Peptide. This result identifies new and unanticipated regions of the TCR to target in the rational design of TCRs for immunotherapy.

### Quaternary Features of the TCR-pMHC

Upon quaternary feature investigation, the LJ-Contacts between the TCR and pMHC dominated bond lifetime prediction. In fact, for all machine learning algorithms investigated, there was no statistically significant i) increase in mean accuracy when expanding to larger feature sets (**Figure 3C**) or ii) decrease in information criteria scores (**Figure 4—supplement 2**). Moreover, although physical features (e.g., *x*-Gyration of TCR) were selected in the exhaustive feature selection process (**Figure 3B**), these did not significantly increase mean accuracy. This demonstrates that no selected physical features improve predictive performance and thus the atomic motion of the TCR or pMHC is unlikely to regulate dissociation kinetics.

### Secondary Features of the TCR-pMHC

To identify the specific subregions of the TCR that determine the TCR-pMHC bond lifetime, we investigated the TCR-pMHC interface and included substructures, or secondary protein features, that defined the interaction (**Figure 1B**). Physiochemical features within each substructure and between adjacent substructures (**Figure 1—supplement 3**) were then evaluated to determine the best predictors of bond lifetime. Among the features and machine learning algorithms selected, a 3-feature combination of secondary features (CDR2β-MHCα(β), CDR1α-Peptide, and CDR3β-Peptide) was selected as the most accurate predictor of TCR-pMHC bond lifetime. This was based on: i) a decrease in information criteria score for 5 of 8 machine learning algorithms; and ii) a statistically significant increase in mean accuracy for 4 of 8 machine learning algorithms when increasing the feature set size from 1 to 3. We found that the combination of total H-bonds between these subregions could predict bond lifetime with the highest accuracy.

The finding that both the total number of unique H-bonds between CDR2β-MHCα(β) and CDR1α-Peptide predict TCR-pMHC bond lifetime is unanticipated. Of particular note, the total number of H-bonds between CDR2β-MHCα(β) remained in all exhaustive search feature sets (**Figure 3B**). Most attention has focused on the heralded CDR3 domains^65^ given the proximity to the peptide (**Figure 1A, B**). In contrast, CDR2 flanks the MHCα(α) and MHCα(β) chains. It is perhaps not surprising, given the significantly larger number of residues (MHCα(β) = 42 residues) compared to the peptide (peptide = 9 residues), that interactions between the CDR2β and the MHCα(β) could potentially be the most significant physiochemical features to impact bond lifetime.

The inclusion of CDR1α-Peptide H-bonds draws new attention to the CDR1α region. Similar to the CDR3β region, CDR1α is in proximity to the peptide (**Figure 1A, B**) and thus hydrogen bonding between these substructures may be expected. However, surprisingly, CDR1α-Peptide H-bonds was exhaustively selected despite interactions between CDR3α-Peptide in the exhaustive feature set (**Figure 4A**). Overall, these results suggest that mutagenesis strategies to increase hydrogen bonding between CDR2β-MHCα(β), CDR1α-Peptide, and CDR3β-Peptide may enhance TCR-pMHC force-dependent bond lifetime. It is important to acknowledge that the interactions between these interfacial substructures may be specific to the DMF5 TCR and will require further investigation to generalize. Nonetheless, these results bring new attention to the CDR1 and CDR2 regions in the future of TCR design. Finally, in contrast to previous reports^28, 57^, peptide radius of gyration and CDR3α-CDR3β distance were not selected in the top ten predictive features. This is likely due to the artificial pMHC unfolding by pulling from TCR-pMHC termini ^27^ and the lack of diversity in TCR-pMHC pairs evaluated ^28, 57^, respectively.

### Computational Methods

One of the limiting factors of this study is the computational constraint of generating a SMD dataset; here, we examined 17 TCR-pMHC pairs. Larger datasets would likely provide more useful insight into feature combinations that predict TCR-pMHC bond lifetime, but come at a significant additional computational cost. Similarly, although the two-layer Elastic Net – Exhaustive Search feature selection methodology provided a rapid filtering of physiochemical features, this biases the machine learning predictor towards features selected by Elastic Net. At the cost of computation, exhaustive or recursive feature selection for each machine learning predictor may improve predictive performance. However, the focus of this work is to provide an architecture for identifying physiochemical features that dictate TCR-pMHC dissociation kinetics.

### Bond Lifetime

The force dependent bond lifetime (at ∼10-20 pN) has been reported to correlate with TCR-pMHC immunogenicity. These findings highlight the importance of TCR-pMHC bond lifetime and suggest that the TCR needs to sustain and form transient bonds under load for sufficient time to initiate biochemical signaling. Thus, we utilized force-dependent bond lifetime as an objective function to uncover the physiochemical determinants of this biomolecular design feature. It is important to note that this biomolecular design feature does not necessarily conflict with catch-slip bond behavior ^24^, and we recognize that our approach may be expanded in the future to include other physiochemical characteristics of the TCR-pMHC bond.

## Conclusion*s*

We have demonstrated the utility of combining two computational methods – steered molecular dynamics and machine learning – to create a methodology that can potentially be used to rapidly and efficiently examine the vast TCR space to predict the bond lifetime, and thus the immunogenic response, of a given TCR-pMHC pair. Our initial results suggest that the physiochemical features of three subregions of the TCR-pMHC are of particular importance in determining bond lifetime (CDR2β-MHCα(β), CDR1α-Peptide, and CDR3β-Peptide) and provide new and unanticipated regions of the TCR to manipulate in the rational design of TCR-engineered T cells.

## SUPPORTING INFORMATION

**Figure 3—supplement 1** Quaternary Features vs Bond Lifetime. **Figure 3—supplement 2** Akaike and Bayesian Information Criterion for Quaternary Features. **Figure 4—supplement 1** Secondary Features vs Bond Lifetime. **Figure 4—supplement 2** Akaike and Bayesian Information Criterion for Secondary Features. **Figure 1—supplement 1** Peptides used in SMD simulations, including their amino acid sequences. **Figure 1—supplement 2** Quaternary Features. **Figure 1—supplement 3** Secondary Features. **Figure 3— supplement 3** Pearson Correlation and Spearman Rank Correlation Coefficients. **Figure 3—supplement 4** Best Machine Learning Models after Hyperparameter Optimization for Quaternary Features. **Figure 4—supplement 3** Best Machine Learning Models after Hyperparameter Optimization for Secondary Features.

## ACKNOWLEDGEMENTS

Simulations were performed on the hpc1/hpc2 clusters in the UC Davis, College of Engineering. This work was supported in part by startup funding to SCG from the Department of Biomedical Engineering. We thank Nuala Del Piccolo for editing the manuscript.

## AUTHOR CONTRIBUTIONS

ZAR: conceptualization, methodology, software, validation, formal analysis, investigation, data curation, writing—original draft, writing—review & editing, visualization

JH: writing—review & editing

IT: methodology, software, validation, formal analysis, writing—review & editing, supervision

RF: conceptualization, methodology, formal analysis, resources, writing—review & editing, supervision, project administration

SCG: conceptualization, methodology, formal analysis, resources, writing—original draft, writing—review & editing, supervision, project administration, funding acquisition

## DATA AVAILABILITY

The datasets of physiochemical features and bond lifetime for each TCR-pMHC pair have been made available on a Dryad repository (https://doi.org/10.25338/B8R33G). Moreover, the root mean square deviation equilibration plots of all TCR-pMHC pairs, starting configurations for all Steered Molecular Dynamics runs, and results from hyperparameter search/cross-validation of the machine learning models have been uploaded to the Dryad repository. The trajectory data generated from Steered Molecular Dynamics simulations are not publicly available due to storage capacity limitations. However, the trajectory data can be reproduced from the initial configurations located in the Dryad repository and are available upon reasonable request, including provision of external storage capacity, to the corresponding author. All scripts including the two-layer Elastic Net – Exhaustive Search algorithm, hyperparameter search of machine learning algorithms with cross validation, and those used to generate figures have been uploaded to a GitHub repository (https://github.com/zrollins/TCR.ai.git). There are no restrictions on data accessibility.

## Supporting Information

**Figure 3—supplement 1:**
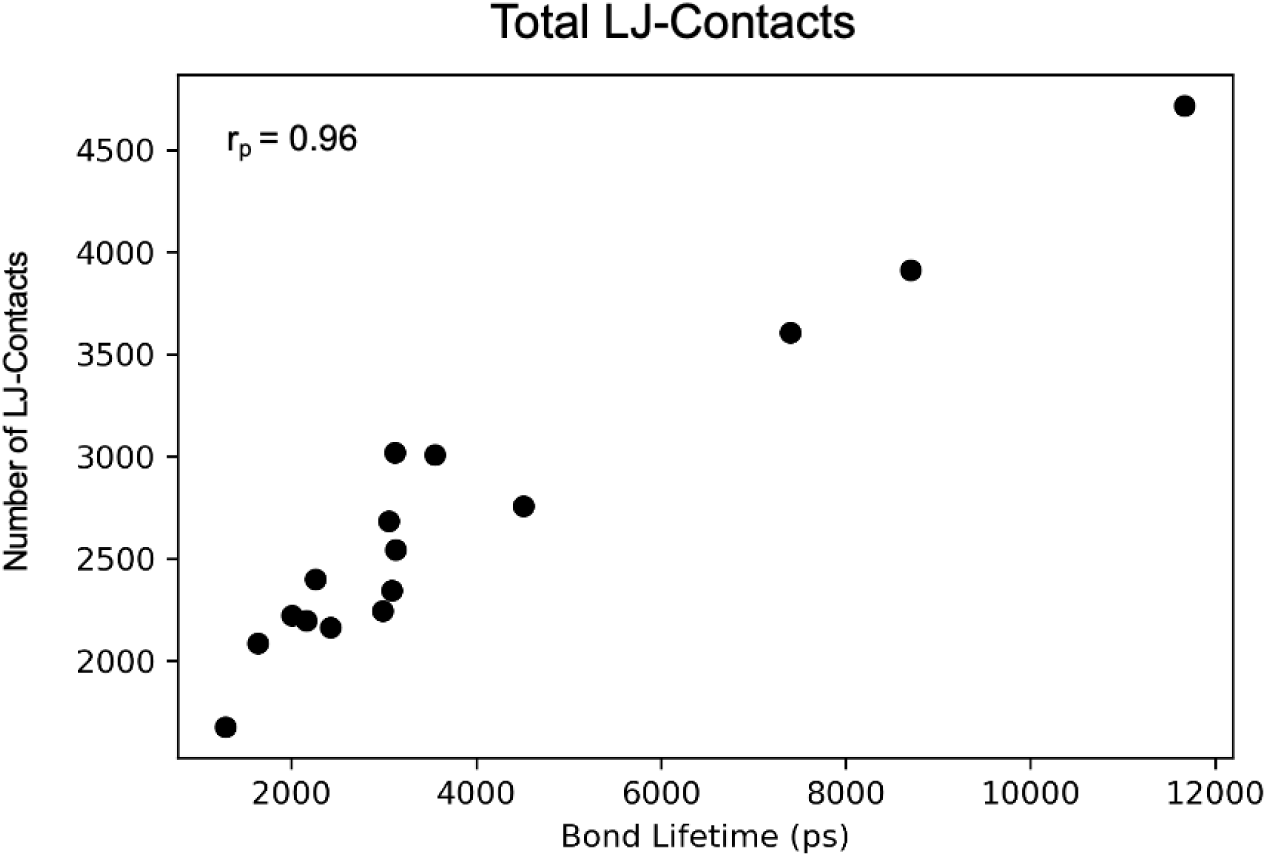
Quaternary Features vs Bond Lifetime. Scatter plot of the total number of LJ-contacts vs bond lifetime for all TCR-pMHC pairs; the Pearson correlation coefficient is listed in the top left corner.

**Figure 3—supplement 2:**
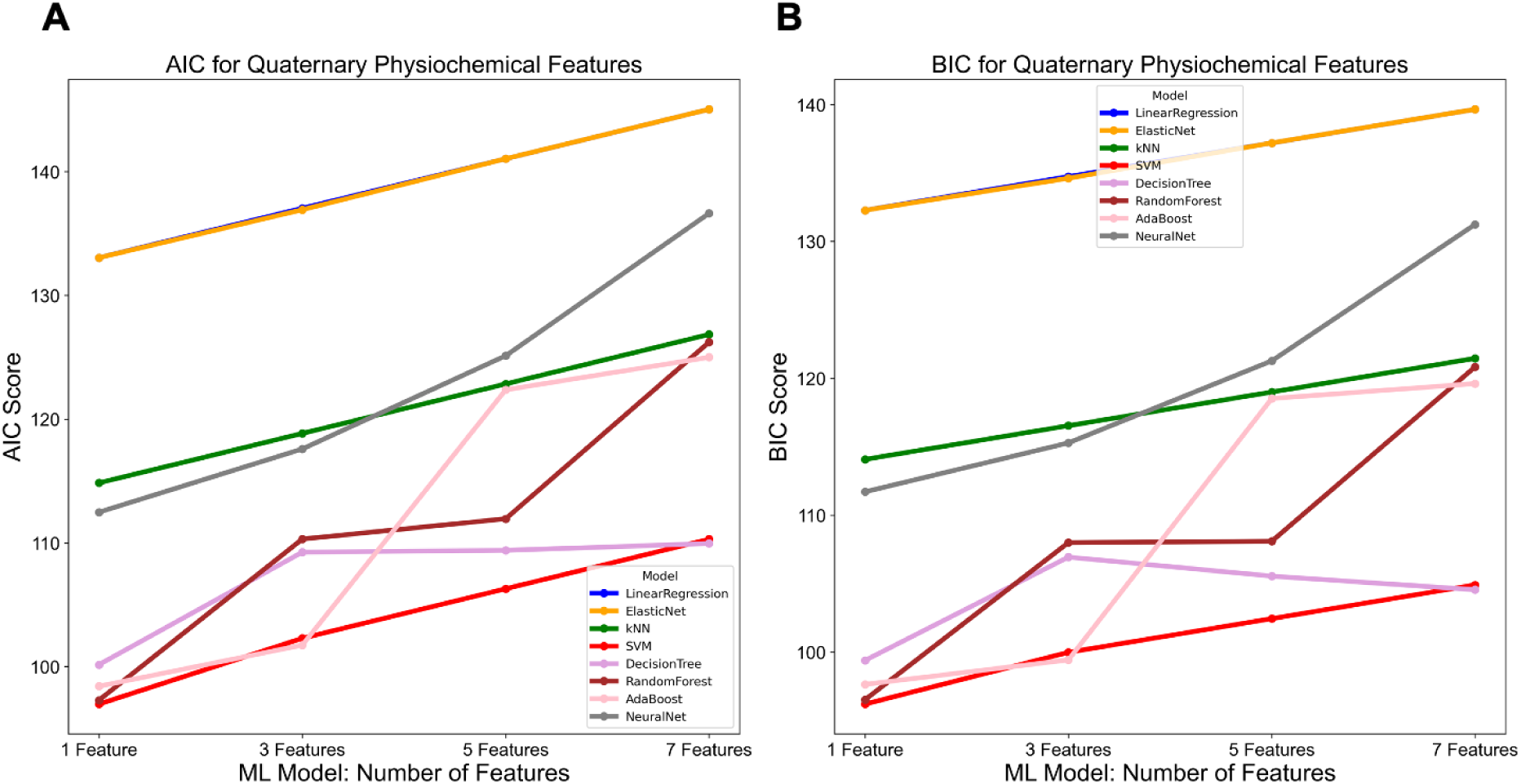
Akaike and Bayesian Information Criterion Scores for Quaternary Features. The **(A)** Akaike Information Criterion (AIC) and **(B)** Bayesian Information Criterion (BIC) for all the quaternary feature sets (i.e., p = 1, 3, 5, and 7) and machine learning models (Linear Regression = blue, Elastic Net = orange, k-Nearest Neighbors = green, Support Vector Machines = red, Decision Tree = purple, Random Forest = brown, AdaBoost = pink, Neural Net = gray).

**Figure 4—supplement 1:**
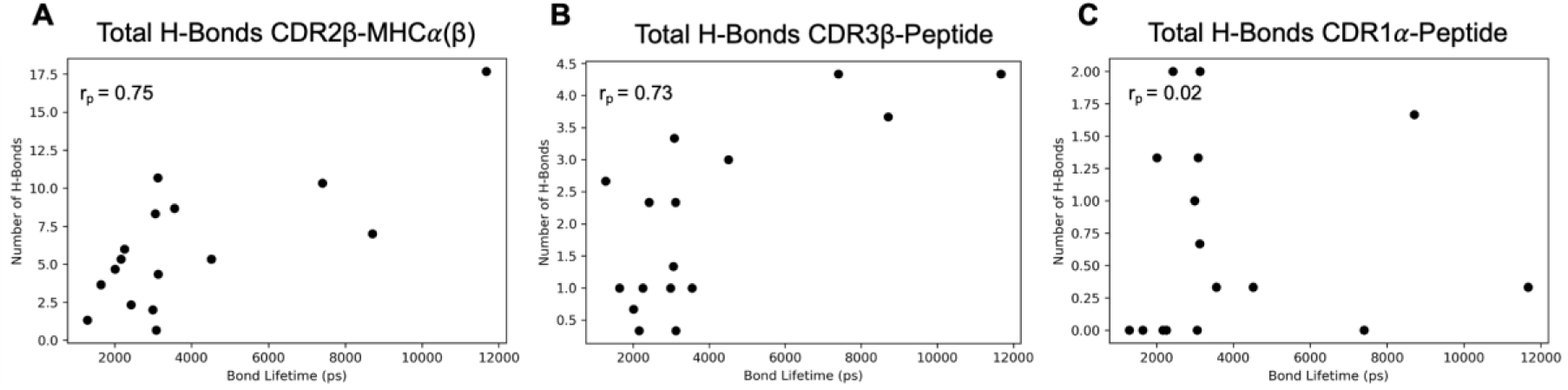
Secondary Features vs Bond Lifetime. Scatter plots of total H-bonds for **(A)** CDR2β -MHCα(β), **(B)** CDR3β-Peptide, and **(C)** CDR1α-Peptide vs Bond Lifetime for all TCR-pMHC pairs; the Pearson correlation coefficient is listed in the top left corner.

**Figure 4—supplement 2:**
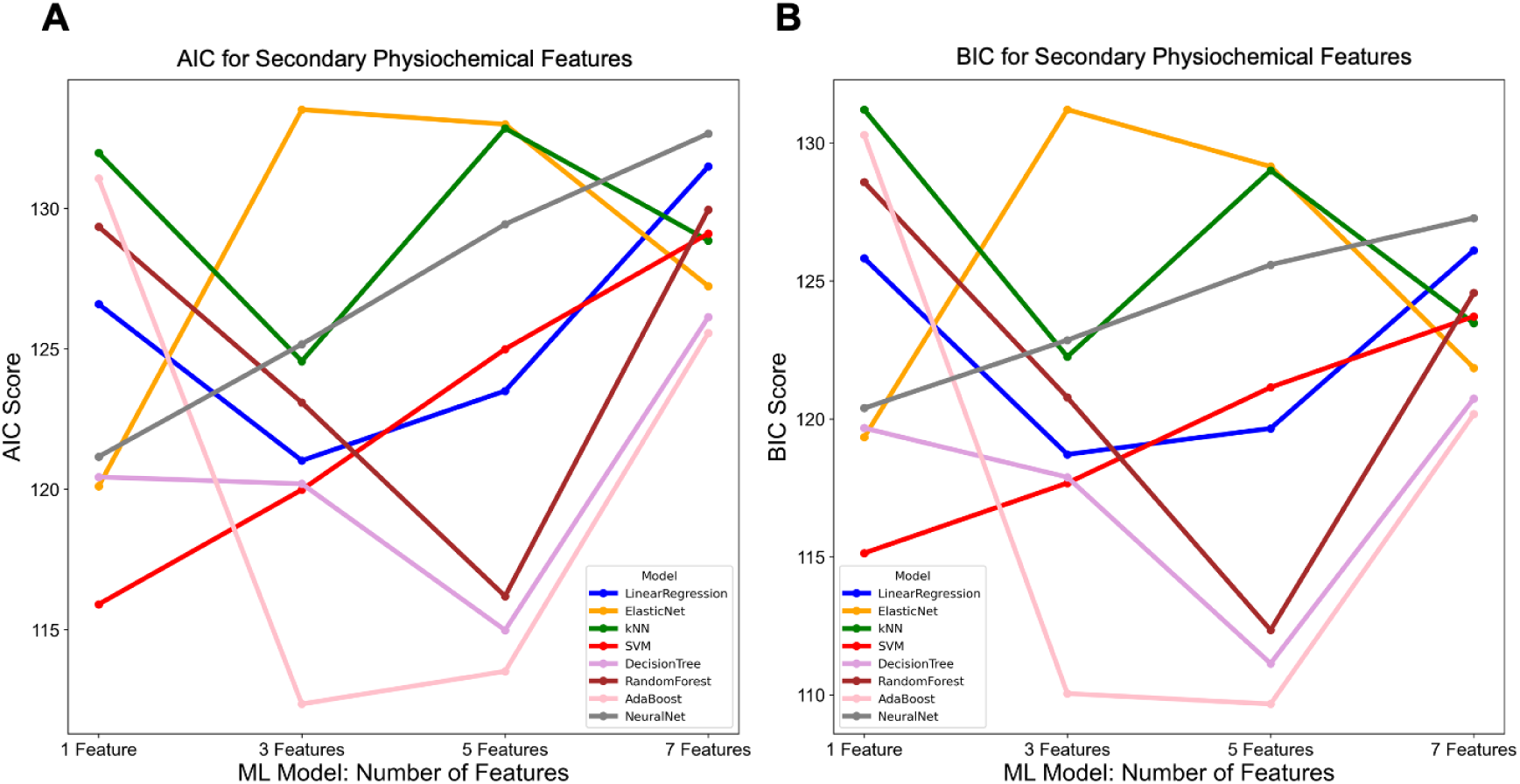
Akaike and Bayesian Information Criterion Scores for Secondary Features. **(A)** The Akaike Information Criterion (AIC) and **(B)** Bayesian Information Criterion (BIC) scores for all secondary feature sets (i.e., p = 1, 3, 5, and 7) and machine learning models (Linear Regression = blue, Elastic Net = orange, k-Nearest Neighbors = green, Support Vector Machines = red, Decision Tree = purple, Random Forest = brown, AdaBoost = pink, Neural Net = gray).

**Figure 1—supplement 1:**
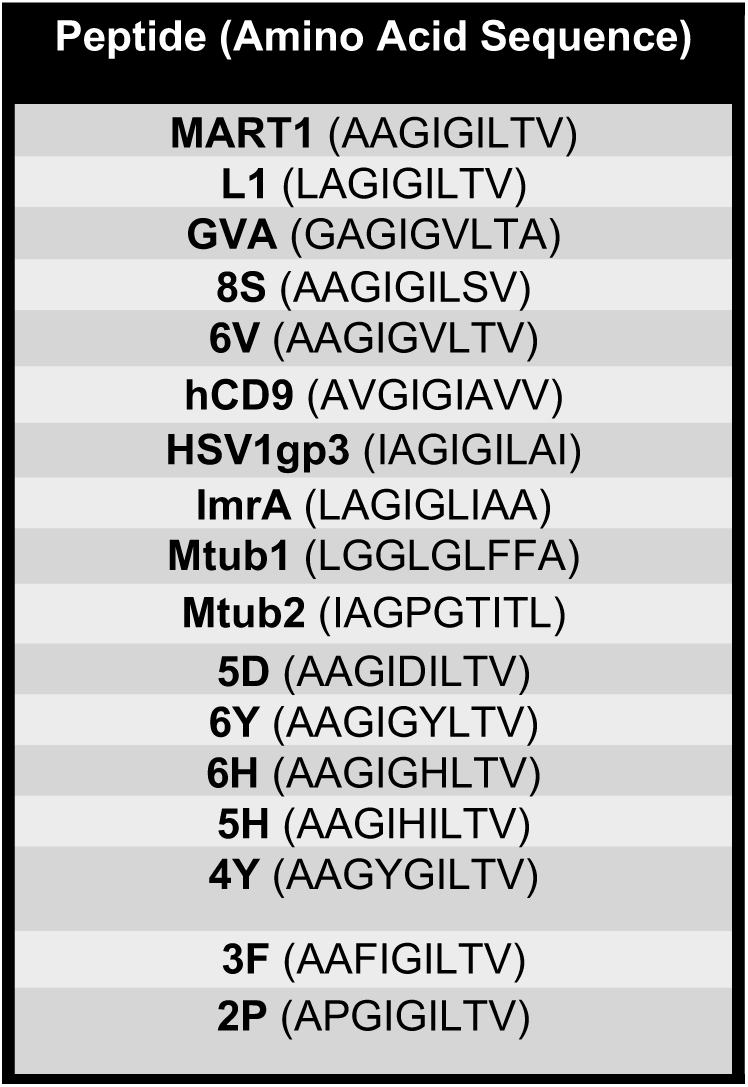
Peptides used in SMD simulations, including their amino acid sequences.

**Figure 1—supplement 2:**
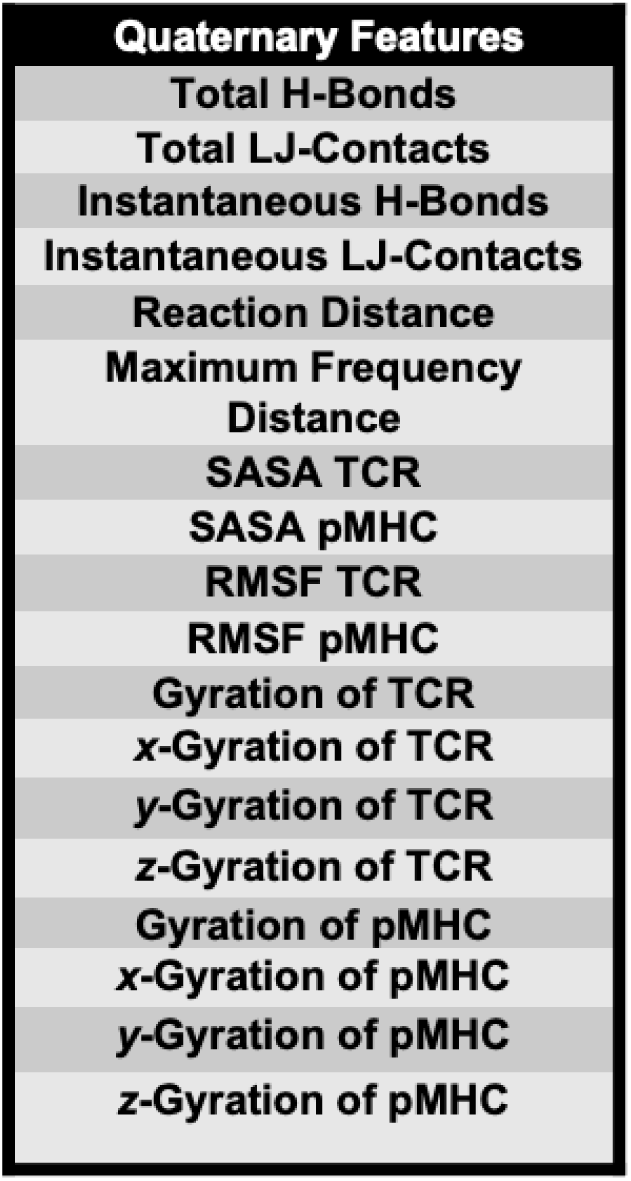
Quaternary Features.

**Figure 1—supplement 3:**
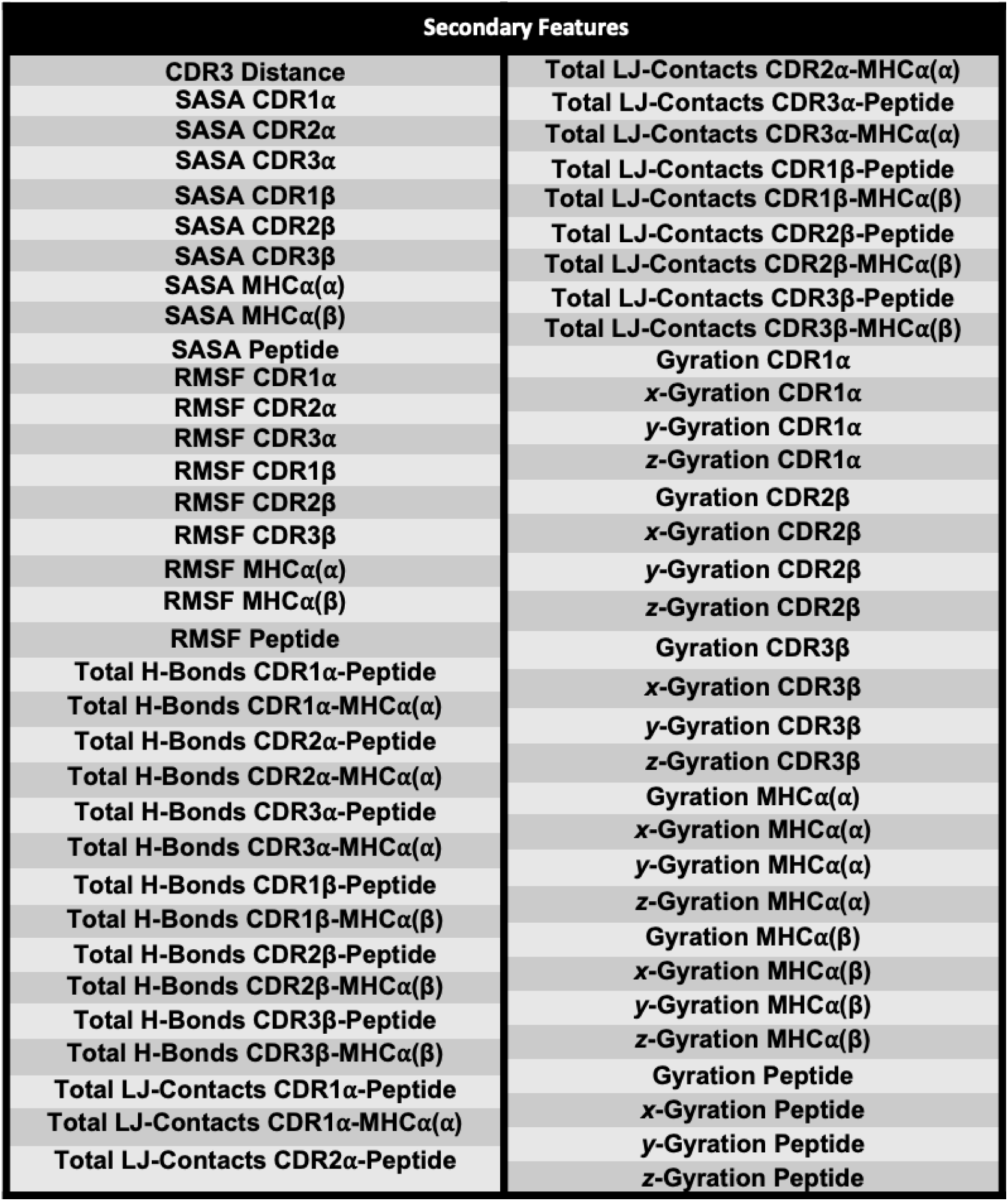
Secondary Features.

**Figure 3—supplement 3:**
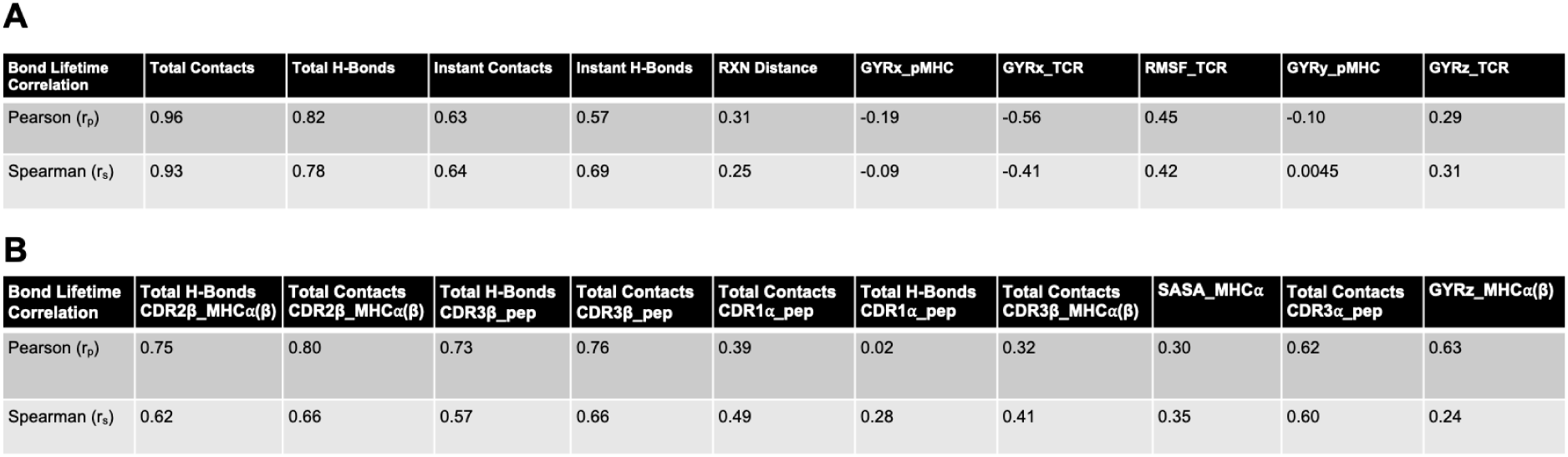
Pearson correlation and Spearman rank correlation coefficients. This includes correlation coefficients for the list of top ten Quaternary Features **(A)** and the list of top ten Secondary Features **(B)**.

**Figure 3—supplement 4:**
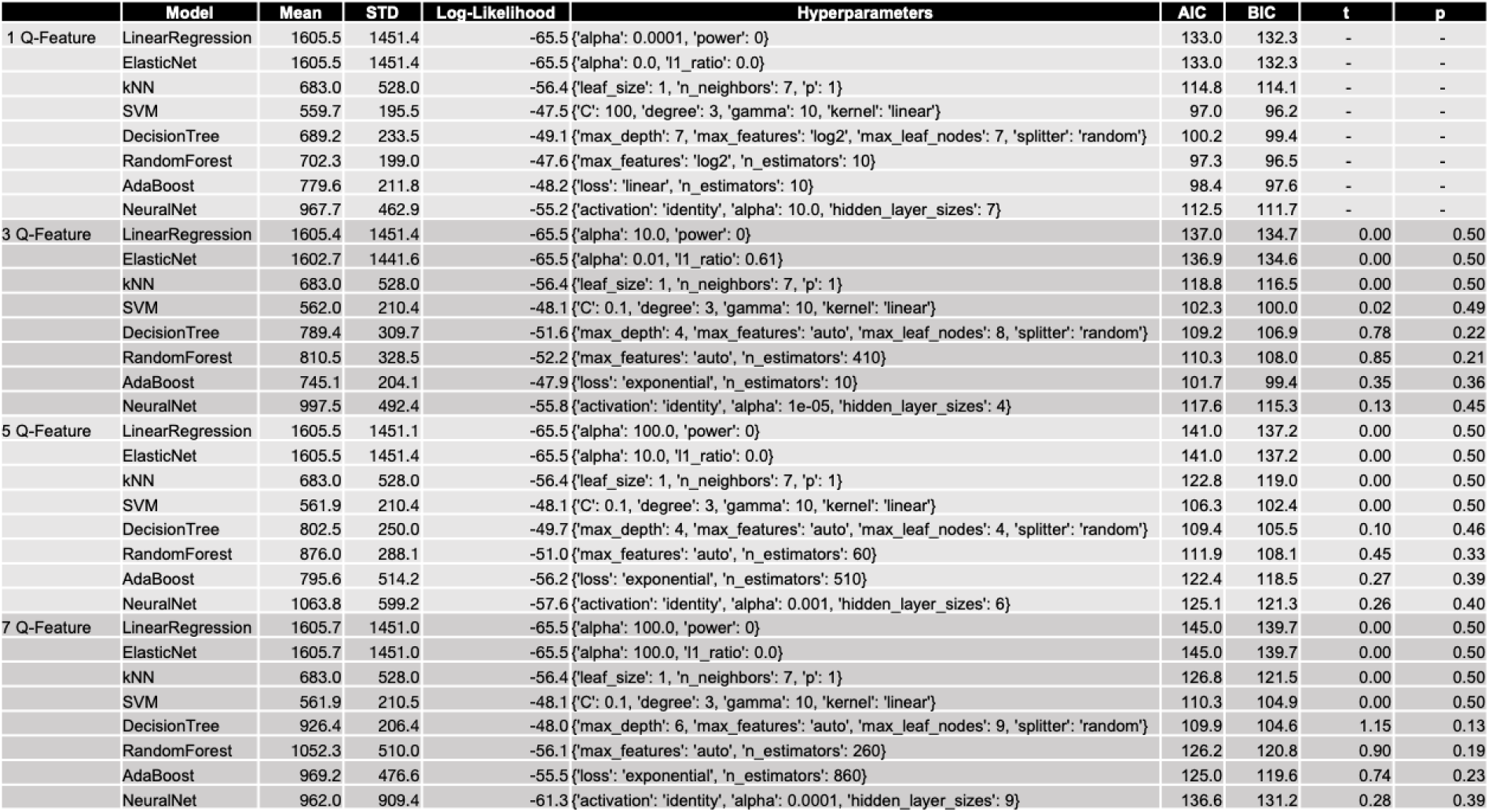
Best Machine Learning Models after Hyperparameter Optimization for Quaternary Features. Table includes the best performing model hyperparameters as well as the mean and standard deviation from repeated threefold cross validation. Akaike and Bayesian Information Criterion are calculated for each model and feature set, based on mean absolute error standard deviation from repeated threefold cross validation, to assess the improved accuracy with increasing complexity. The respective algorithms are statistically compared across feature sets using a one-tailed student’s t-test.

**Figure 4—supplement 3:**
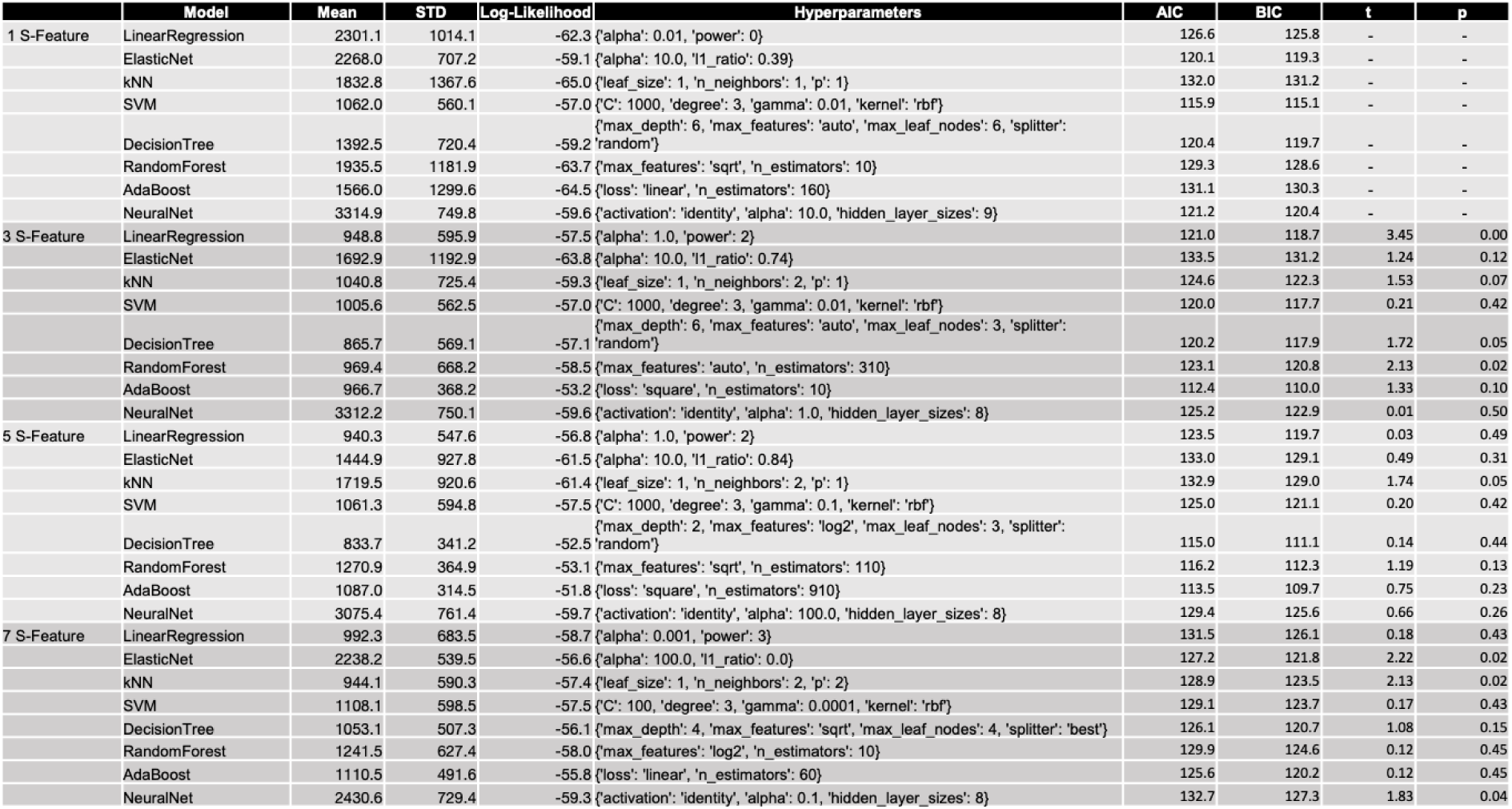
Best Machine Learning Models after Hyperparameter Optimization for Secondary Features. Table includes the best performing model hyperparameters as well as the mean and standard deviation from repeated threefold cross validation. Akaike and Bayesian Information Criterion are calculated for each model and feature set, based on mean absolute error standard deviation from repeated threefold cross validation, to assess the improved accuracy with increasing complexity. The respective algorithms are statistically compared across feature sets using a one-tailed student’s t-test.

